# Pathway incompatibility between NF-κB and RAS signaling constrains oncogenicity in B-cell leukemia

**DOI:** 10.64898/2026.03.04.709574

**Authors:** Emma Stewart, Mia Knupke, Adriana Collins, Hua Lou, Lai N. Chan

## Abstract

B-cell acute lymphoblastic leukemia (B-ALL) arises from lymphoid precursors that fail to complete normal B-cell maturation and frequently harbor activating mutations in the RAS-ERK pathway that promote leukemic survival. These cells remain dependent on precursor B-cell receptor (pre-BCR)-associated signaling programs, which oncogenic RAS can functionally mimic. While oncogenesis is often attributed to cooperating mutations, mutually exclusive genetic alterations suggest that certain oncogenic pathways may be functionally incompatible. In our previous work, we proposed the concept of pathway incompatibility, in which co-activation of specific oncogenic signaling programs antagonizes their oncogenic activity and suppresses tumorigenesis.

Here, we examined the interaction between RAS signaling and canonical NF-kB activity in B-ALL within this framework. Genomic analysis revealed underrepresentation of cases harboring concurrent RAS and NF-kB pathway alterations. Functionally, activation of canonical NF-kB induced apoptotic depletion of RAS-pathway-mutated B-ALL cells, whereas activation of the non-canonical pathway conferred a growth advantage. Mechanistically, canonical NF-kB suppressed pre-BCR-associated signaling programs and increased expression of mature B-cell receptor (BCR) components, indicating a developmental signaling state incompatible with oncogenic RAS activity.

Consistent with this framework, enforced oncogenic RAS signaling was poorly tolerated in BCR-positive lymphoma cells unless BCR expression was disrupted. Pharmacologic activation of NF-kB reduced ERK signaling and selectively impaired viability of RAS-pathway-mutated B-ALL cells, with enhanced effects in combination with ERK inhibition. Together, these findings identify canonical NF-kB signaling as a developmental context-dependent constraint on RAS-driven leukemogenesis and support pathway incompatibility as a determinant of the oncogenicity of signaling pathways.

## INTRODUCTION

B-cell acute lymphoblastic leukemia (B-ALL) is one of the most common malignancies of childhood, with more than 5,000 new cases diagnosed annually in the United States^1–3^. The disease is characterized by disruption of normal B-cell maturation, leading to accumulation of lymphoid precursors unable to progress through developmental checkpoints^4–5^. During normal B-cell development, early precursors express the precursor B-cell receptor (pre-BCR), composed of an immunoglobulin heavy chain paired with surrogate light chains (SLC; VpreB and λ5), which promotes survival and proliferation of early B-cell precursors. Replacement of surrogate with conventional light chains generates the mature B-cell receptor (BCR), enabling antigen recognition and immune function^6–7^. In B-ALL, leukemic cells are typically arrested prior to this transition, thereby evading differentiation signals while sustaining proliferative capacity^4–5^.

Activating mutations in the RAS–ERK pathway occur in approximately 35% of B-ALL cases^8–11^ and functionally mimic pre-BCR-dependent signaling, sustaining leukemic cell survival and proliferation in the absence of normal B-cell maturation^11-13^. Our prior work showed that activating lesions in the RAS-ERK pathway functionally mimic pre-BCR signaling through induction of BCL6, thereby providing survival and proliferative signals^11^. In support of this scenario, deletion of the pre-BCR adaptor *Blnk* suppresses RAS-driven leukemogenesis^11^. Clinically, RAS pathway activation correlates with therapy resistance, relapse, and central nervous system involvement^8–10^, emphasizing the need for therapeutic strategies that exploit vulnerabilities associated with developmental arrest.

Cancer progression is often conceptualized as driven by cooperating oncogenic alterations^14–15^, and mutually exclusive mutations observed across tumor types are frequently interpreted as reflecting functional redundancy^16–17^. However, evidence from B-ALL suggests an alternative model in which certain oncogenic pathways are mutually exclusive because their signaling outputs are antagonistic rather than redundant. This phenomenon, termed *pathway incompatibility*, arises when co-activation results in reciprocal inhibition and suppression of malignant phenotypes^11^. Building on this framework, our findings suggest that the relationship between RAS signaling and canonical NF-κB activity represents a potential example of pathway incompatibility, providing insight into how developmental signaling context constrains leukemogenesis.

## RESULTS

### Genomic interaction analysis identifies antagonism between NF-κB and RAS pathways in B-ALL

NF-κB signaling exerts context-dependent effects across B-cell malignancies, functioning as either an oncogene or tumor suppressor^18-22^. In Philadelphia chromosome–positive (*Ph*^+^) B-ALL, driven by the BCR-ABL1 fusion kinase, NF-κB activity is essential for malignant transformation; as its inhibition impairs cell growth and enhances tyrosine kinase inhibitor sensitivity^20-21^. Notably, BCR-ABL1 also activates STAT5, which promotes leukemogenesis in part by blocking B-cell differentiation. In this context, NF-κB has been shown to counteract STAT5-driven transcriptional programs^22^. These observations underscore the complexity and context dependence of NF-κB signaling in lymphoid transformation.

To evaluate potential interactions between oncogenic pathways, we conducted mutation co-occurrence analysis across 572 previously reported B-ALL cases^23-26^. Activating mutations affecting RAS-pathway components were detected in 266 cases, whereas NF-κB–pathway activating alterations were identified in 60 cases. Based on mutation frequencies, 28 cases would be expected to harbor both alterations if they occurred independently; however, only 8 such cases were observed (odds ratio = 0.15, *P* < 0.0001), indicating that B-ALL samples carrying both lesions occurred far less frequently than expected by chance. This marked underrepresentation indicates a potential negative genetic interaction between the RAS and NF-κB pathways and raises the possibility of functional antagonism between these signaling programs.

### Canonical NF-κB activation suppresses growth of RAS-pathway mutated B-ALL cells

NF-κB signaling operates through two mechanistically distinct pathways: canonical and non-canonical. Canonical signaling activates the IKK2 (IKKβ)/NEMO complex to release p65:p50 dimers that regulate genes involved immune responses, survival, and proliferation^27-29^. The non-canonical pathway engages NF-κB inducing kinase (NIK, also known as MAP3K14)) and IKK1 (IKKα) to generate p52:RelB complexes that primarily govern lymphocyte development and organogenesis^27-29^. Prior studies demonstrated opposing functions of canonical and non-canonical NF-κB signaling in dendritic cells during radiotherapy, with canonical NF-κB required for radiation-induced anti-tumor immunity, whereas activation of the non-canonical pathway suppresses type I interferon production and limits immune activation^30^.

To determine how NF-κB signaling influences the growth of RAS-pathway–mutated B-ALL cells, we selectively activated each NF-κB cascade in wild-type mouse B-cell precursor cells transformed with NRAS^G12D^, a model used in our prior work to study pathway incompatibility between STAT5- and RAS-signaling in B-ALL^11^. In B-ALL, recurrent RAS-pathway lesions most commonly involve activating mutations in NRAS (39.6%; e.g., NRAS^G12D^)^8^.

Previous studies showed that overexpression of wild-type NIK increases p52:RelB abundance and DNA binding, thereby activating non-canonical NF-κB signaling^31,32^. To compare the effects of non-canonical and canonical NF-κB activation in RAS-driven context, we expressed wild-type NIK or IKBKB S177E/S181E (constitutively active IKK2, Ikk2ca)^33, 34^ in mouse NRAS^G12D^ B-ALL cells. Using this approach, we found that NIK overexpression promoted cell growth (**Fig. 1a**). In contrast, activation of the canonical pathway through Ikk2ca resulted in depletion of cells from culture in growth competition assays (**Fig. 1a**). This depletion was associated with a ∼50% (approximately 1.5-fold) increase in Annexin V-positive, SYTOX Blue-negative cells, consistent with induction of early apoptosis following canonical NF-κB activation (**Fig. 1b**). Consistent with a growth-restrictive role for canonical signaling, shRNA-mediated knockdown of *Nfkb1* conferred a competitive advantage (**Fig. 1c**). The super-repressor form of IkBα (IkBα-SR) is the dominant variant of IkBα and blocks the nuclear translocation of NF-κB. Similar to repression of *Nfkb1*, overexpression of IkBα-SR promoted cell growth of RAS-pathway–mutated B-ALL cells (**Fig. 1d**).

**Figure 1:**
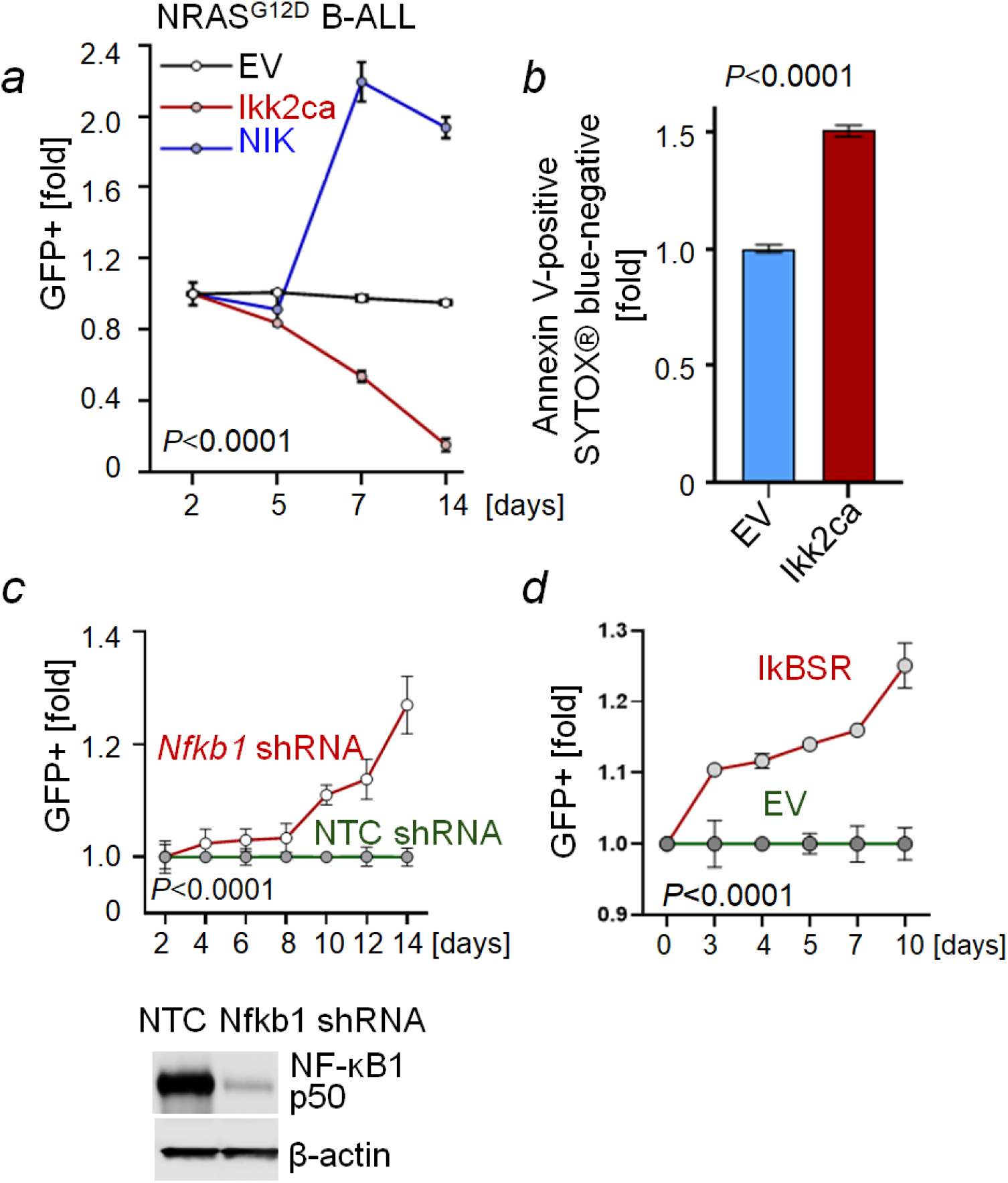
Antagnoism between ERK- and canonical NF-kB activation in RAS-pathway mutated B-ALL. (***a***) Enrichment or depletion of mouse NRAS^G12D^ B-ALL cells transduced with empty vector (EV) control, Ikk2ca-GFP (canonical NF-κB activation), or NIK wild-type-GFP (non-canonical NF-κB activation) as measured by flow cytometry (n=3). (***b***) Following transduction with EV control or Ikk2ca, NRAS^G12D^ B-ALL cells were analyzed by flow cytometry for Annexin V-positive and SYTOX blue stain-negative populations, indicative of early apoptotic cells (n=3). (***c***) Enrichment or depletion of mouse NRAS^G12D^ B-ALL cells transduced with GFP-tagged non-targeting control (NTC) or *Nfkb1*-targeting shRNA, as measured by flow cytometry (n=3; top). Western blots of NRAS^G12D^ B-ALL cells expressing NTC shRNA or *NFKB1*-targeting shRNA (bottom). (***d***) Enrichment or depletion of NRAS^G12D^ B-ALL cells transduced with empty vector (EV) control or GFP-tagged IkBα super-repressor (IkBSR), n=3, as measured by flow cytometry.

We next examined whether responses to canonical NF-κB activation depended on Erk signaling activity. To do this, we compared wild-type B-cell precursor cells with *Nf1 23aIN/23aIN* cells, a mutant mouse model in which altered splicing of *Nf1* reduces neurofibromin Ras-GAP activity and selectively enhances RAS–ERK signaling without significantly affecting PI3K–AKT signaling^**35**^. Consistent with prior studies, *Nf1 23aIN/23aIN* B-cell precursor cells exhibited elevated Erk1/2 phosphorylation relative to wild-type cells (**Fig. 2a**). In growth competition assays, cells with higher Erk activity (*Nf1 23aIN/23aIN*) showed markedly greater depletion following Ikk2ca expression compared with cells displaying lower Erk activity (wild-type) (**Fig. 2b**). These findings indicate that the growth-suppressive effect of canonical NF-κB is more pronounced in cells with elevated Erk signaling.

**Figure 2:**
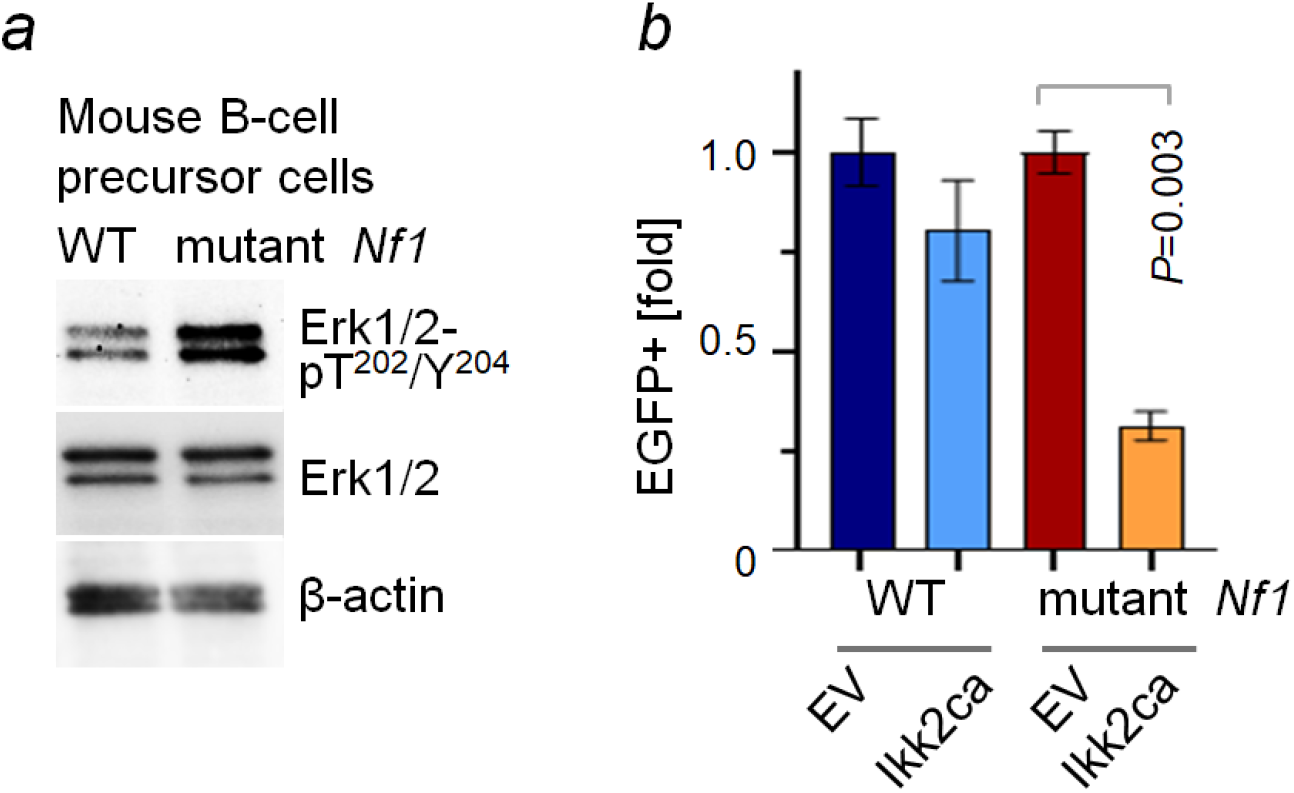
Canonical NF-κB–mediated growth suppression depends on ERK signaling level. (a)Western blots of mouse B-cell precursor cells showing low and high levels of Erk1/2 phosphorylation in wild-type (WT) and *Nf1 23aIN/23aIN* (*Nf1* mutant) B-cell precursor cells, respectively. (***b***) Relative enrichment or depletion of WT (low Erk activity) and *Nf1 23aIN/23aIN* (*Nf1* mutant; high Erk activity) mouse B-cell precursor cells following transduction with empty vector (EV) control or Ikk2 ca-GFP (canonical NF-κB activation), measured by flow cytometry (n=3).

Western blot analyses further supported antagonism between these pathways. NRAS^G12D^ expression in mouse B-cell precursor cells increased ERK1/2 phosphorylation while reducing phosphorylation of NF-κB p65 (**Fig. 3a**). Conversely, activation of canonical NF-κB through Ikk2ca enhanced p65 phosphorylation and was accompanied by decreased ERK1/2 phosphorylation (**Fig. 3b**). Together, these results demonstrate opposing effects of canonical NF-κB and RAS–ERK signaling on pathway activity. Because these responses were specific to canonical activation, subsequent analyses focus on this branch of the pathway, hereafter referred to as NF-κB.

**Figure 3:**
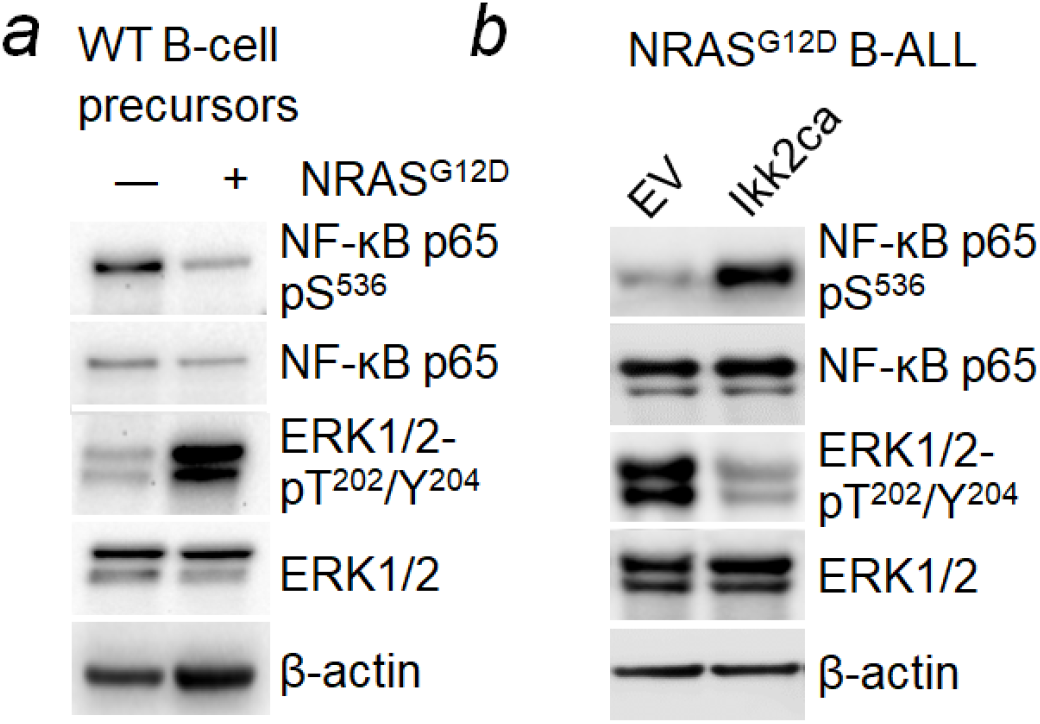
Canonical NF-κB and oncogenic RAS display reciprocal signaling antagonism. (a)Western blots of wild-type (WT) mouse B-cell precursor cells with or without NRAS^G12D^. (***b***) Western blots of NRAS^G12D^ B-ALL cells expressing empty vector (EV) control or Ikk2ca.

### Canonical NF-κB alters receptor composition toward a BCR-associated state

Notably, activation of canonical NF-κB through Ikk2ca reduced phosphorylation of the receptor-proximal kinase Syk (Tyr352), indicating attenuation of pre-BCR–associated signaling (**Fig. 4a**). Consistent with disruption of early B-cell signaling programs, NF-κB activation increased phosphorylation of Foxo1 at S256 (**Fig. 4a**), indicative of Foxo1 inactivation. In line with these changes, Bcl6 expression was decreased (**Fig. 4a**), a downstream effector of RAS signaling that functionally mimics pre-BCR activity to sustain survival and proliferation of RAS-driven B-ALL cells^11^. Together, these observations indicate that canonical NF-κB signaling suppresses both receptor-proximal signaling and downstream transcriptional programs required for maintenance of the pre-BCR state.

**Figure 4:**
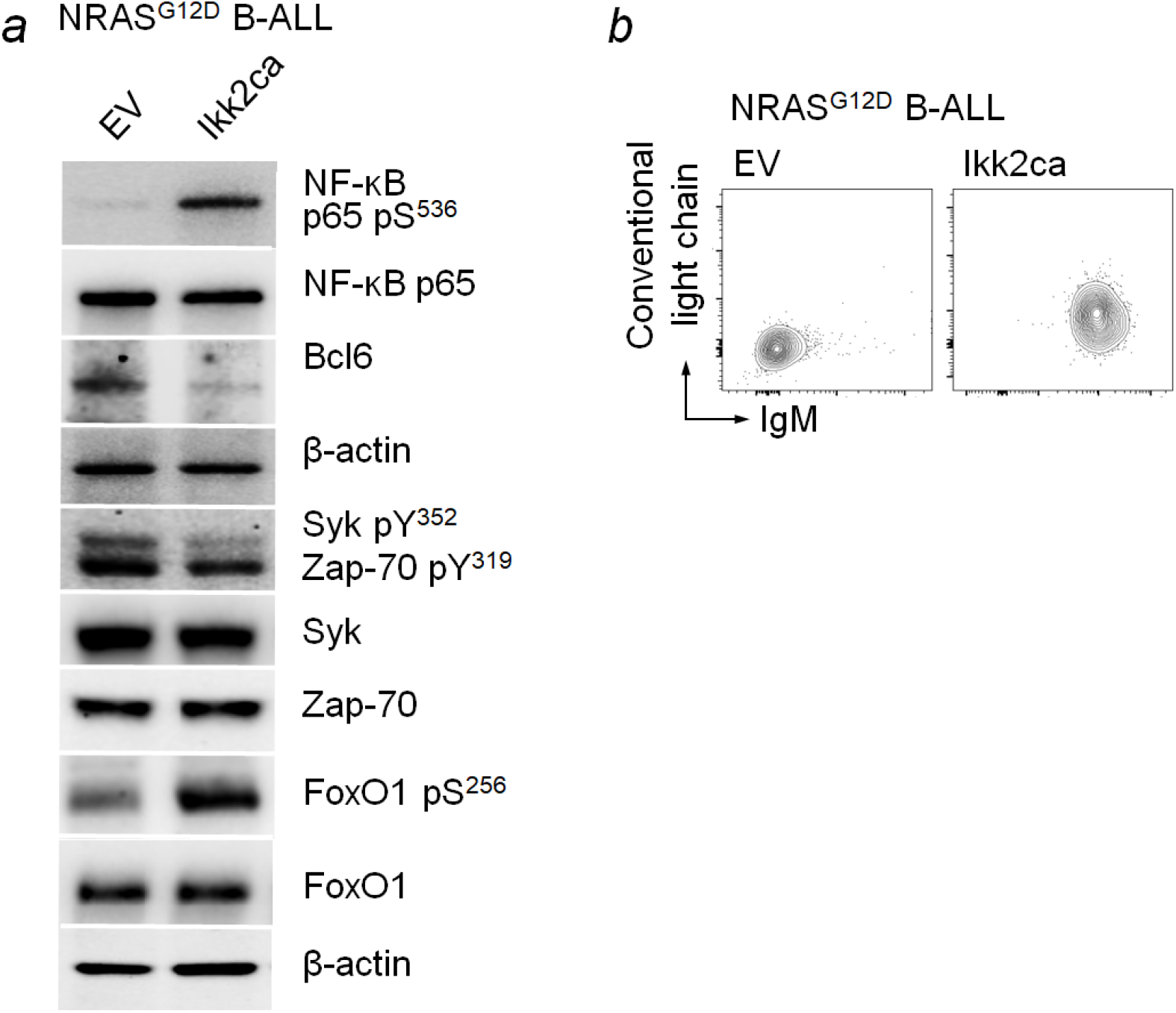
Canonical NF-κB suppresses pre-BCR programs and promotes acquisition of BCR components. (***a***) Western blots of NRAS^G12D^ B-ALL cells expressing empty vector (EV) control or Ikk2ca. (***b***) Flow cytometry analysis of BCR components (conventional λ light chains and IgM) on mouse NRAS^G12D^ B-ALL cells upon overexpression of empty vector (EV) control or Ikk2ca.

During normal B-cell development, SLCs are replaced by conventional κ or λ light chains (cLCs), which pair with μ heavy chains to form the mature B-cell receptor (BCR), enabling B-cell differentiation^36-39^. Although leukemia blasts are typically arrested prior to BCR expression^4,5^, activation of NF-κB increased surface expression of cLCs and IgM (μ heavy chain) (**Fig. 4b**). Because coordinated expression and trafficking of both chains to the plasma membrane are required for receptor assembly, these changes indicate enhanced formation of surface BCR complexes.

Together, these findings suggest that canonical NF-κB activation shifts receptor signaling away from pre-BCR– associated programs and toward a BCR-like state, consistent with partial developmental progression that is incompatible with RAS-driven leukemogenesis.

### Developmental signaling context constrains oncogenic RAS signaling

Because activation of canonical NF-κB increased expression of mature BCR components and shifted leukemic cells away from pre-BCR–associated signaling programs, we next examined whether this change in signaling context might contribute to the reduced tolerance of oncogenic RAS activity. Activating RAS-pathway mutations were enriched in malignancies arising from early or terminal stages of B-cell development lacking surface BCR expression, including B-ALL and multiple myeloma, but were uncommon in tumors derived from mature BCR-expressing B cells^40^. Consistent with this pattern, oncogenic RAS efficiently transforms immature B-cell precursors in murine models^11^ yet fails to transform mature BCR^+^ B cells^41,42^. These observations suggest that developmental signaling context may influence whether oncogenic RAS can support leukemic growth.

To directly assess whether oncogenic RAS is incompatible with the BCR-positive state, we expressed empty vector (EV) control or NRAS^G12D^ in diffuse large B-cell lymphoma (DLBCL) cells that retain surface expression of BCR component conventional κ light chains (**Fig. 5a**). Expression of NRAS^G12D^ resulted in depletion of cells in growth competition assays (**Fig. 5b**), indicating that oncogenic RAS signaling is poorly tolerated in these cells.

**Figure 5:**
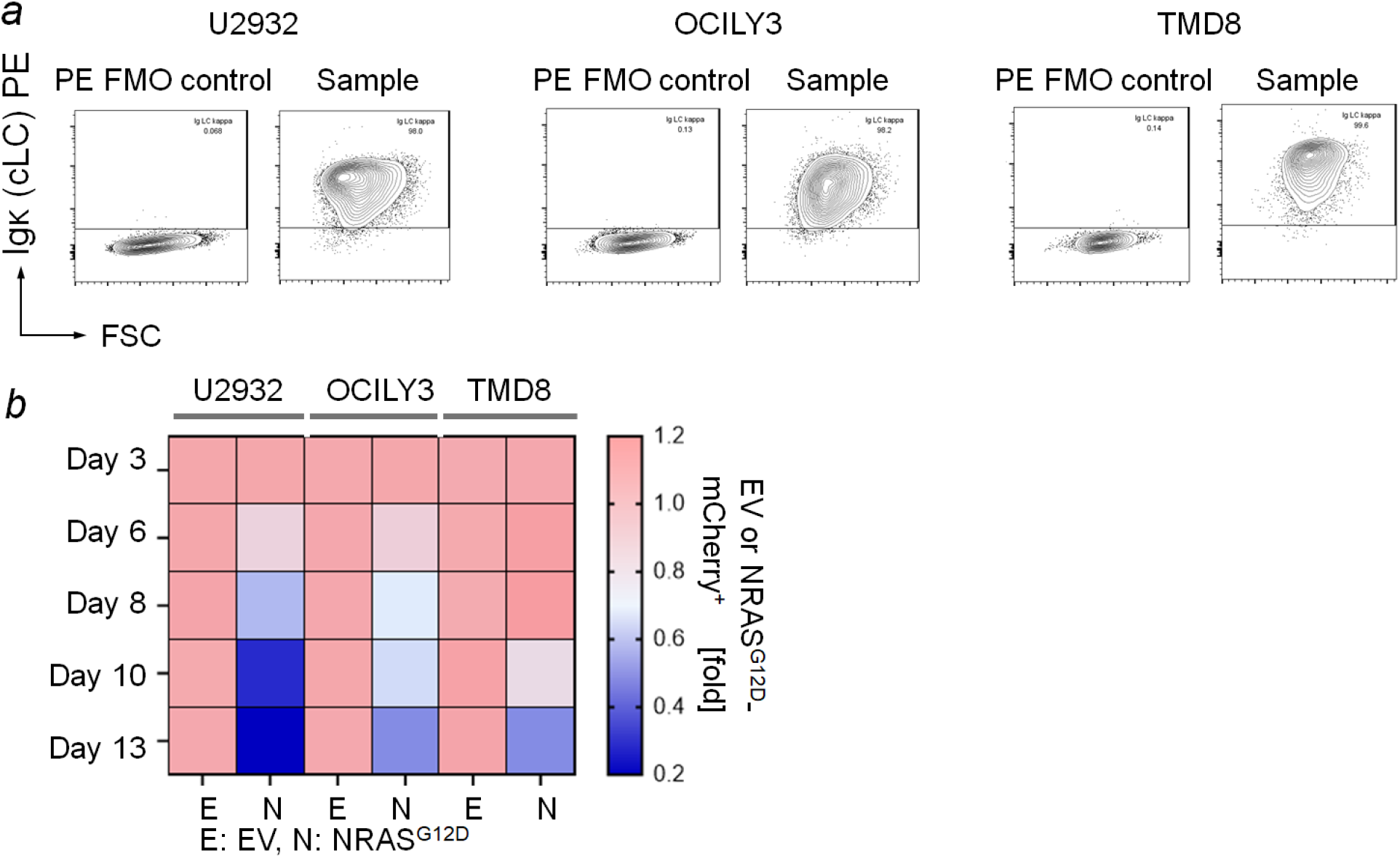
Expression of mutant NRAS depletes BCR-positive lymphoma cells. (a)Flow cytometry analysis of conventional light chains (Ig kappa LC) on human diffuse large B-cell lymphoma (DLBCL) cells. Surface expression of Ig kappa LC was shown on viable cells (SYTOX® blue stain-negative). FMO: Fluorescence Minus One (cells were stained with SYTOX® blue stain but not PE-conjugated antibodies against Ig kappa LC. (***b***) Cells were transfected with plasmid expressing mCherry-tagged empty vector (EV) control or NRAS^G12D^ using electroporation. Following transfection, enrichment or depletion of mCherry^+^ cells was monitored by flow cytometry. Shown are average fold changes (compared to day 3 post transfection) from 3 independent replicates.

To determine whether this observed depletion of NRAS^G12D^-expressing cells was linked to BCR expression, we used CRISPR/Cas9 to disrupt the constant region of conventional κ light chains in U2932 cells (**Fig. 6**). In growth competition assays, NRAS^G12D^ expression reduced the abundance of cells retaining conventional light-chain expression (**Fig. 6b**), whereas loss of conventional light chains markedly rescued this depletion (*P* = 0.0078; **Fig. 6a**). Together, these findings indicate that the signaling environment associated with BCR expression constrains the oncogenic activity of RAS and suggest that the NF-κB–induced shift toward a BCR-like state may contribute to suppression of RAS-driven leukemic programs.

**Figure 6:**
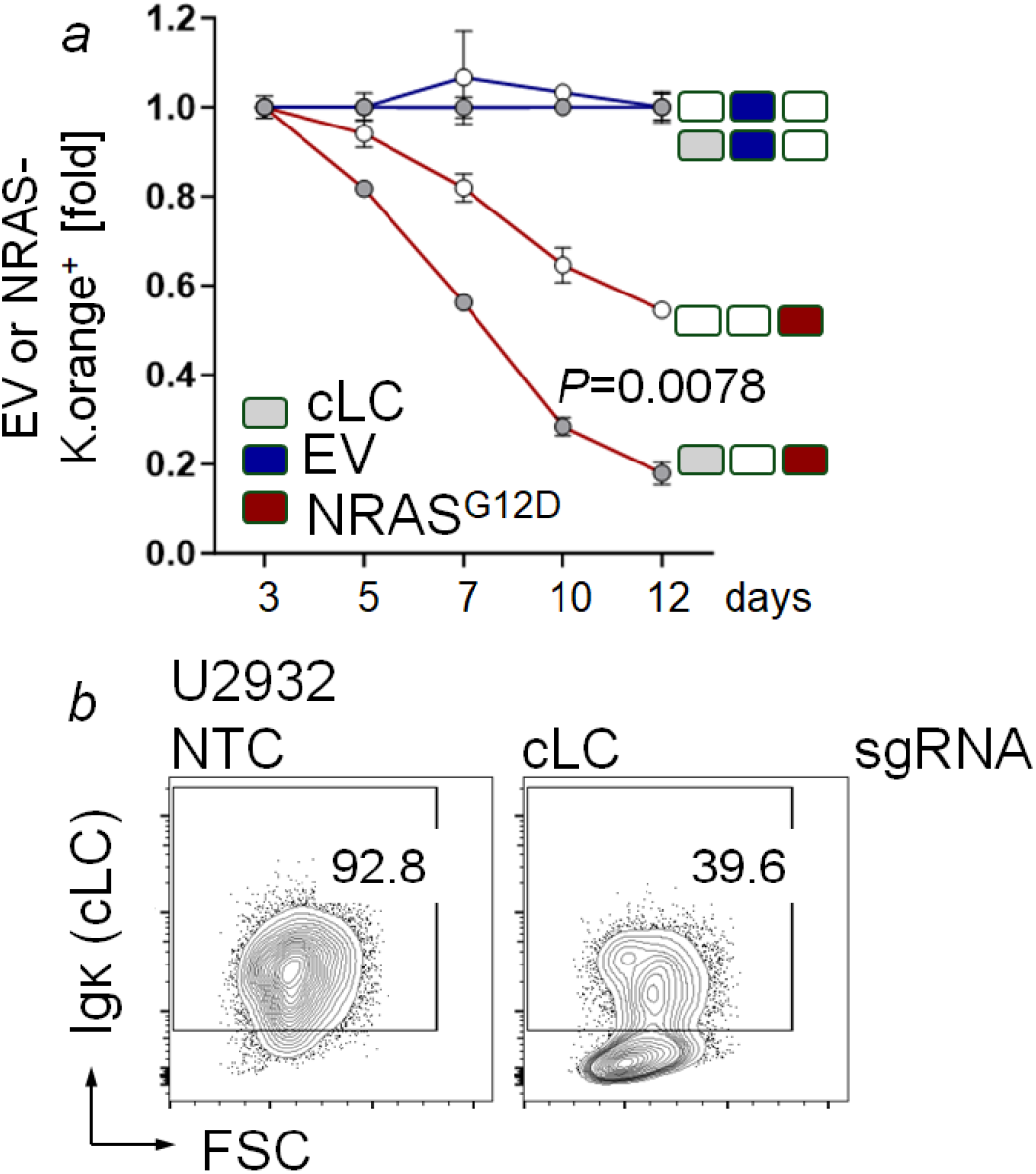
BCR expression mediates intolerance to oncogenic RAS signaling. (***a-b***) U2932 cells were electroporated with Cas9 ribonucleoprotein (RNPs) complexes containing recombinant Cas9 and either non-targeting control (NTC) or conventional light chain (cLC; *IGKC*)-targeting guide RNAs (***b***). Following transfection, cells were sorted into cLC-positive or cLC-negative populations from NTC or cLC sgRNA-transfected cells, respectively. Sorted cells were transfected with EV or NRAS^G12D^-K.orange using electroporation. Enrichment or depletion of mCherry^+^ cells was monitored by flow cytometry (n=3) (***a***). Gray box: cLC expressed. **Blue** box: EV expressed. **Red** box: NRAS^G12D^ expressed.

### Antagonistic interaction between RAS and NF-κB signaling reveals potential therapeutic relevance

We next examined whether the functional opposition between NF-κB and RAS signaling could be leveraged in a therapeutic context. Across a panel of human B-ALL cell lines^43,44^, responses to canonical NF-κB activation varied according to baseline ERK activity. Cell lines with the highest ERK signaling were preferentially depleted in growth competition assays following IKK2ca expression, whereas cell lines with lower ERK activity exhibited relative enrichment (**Fig. 7**).

**Figure 7:**
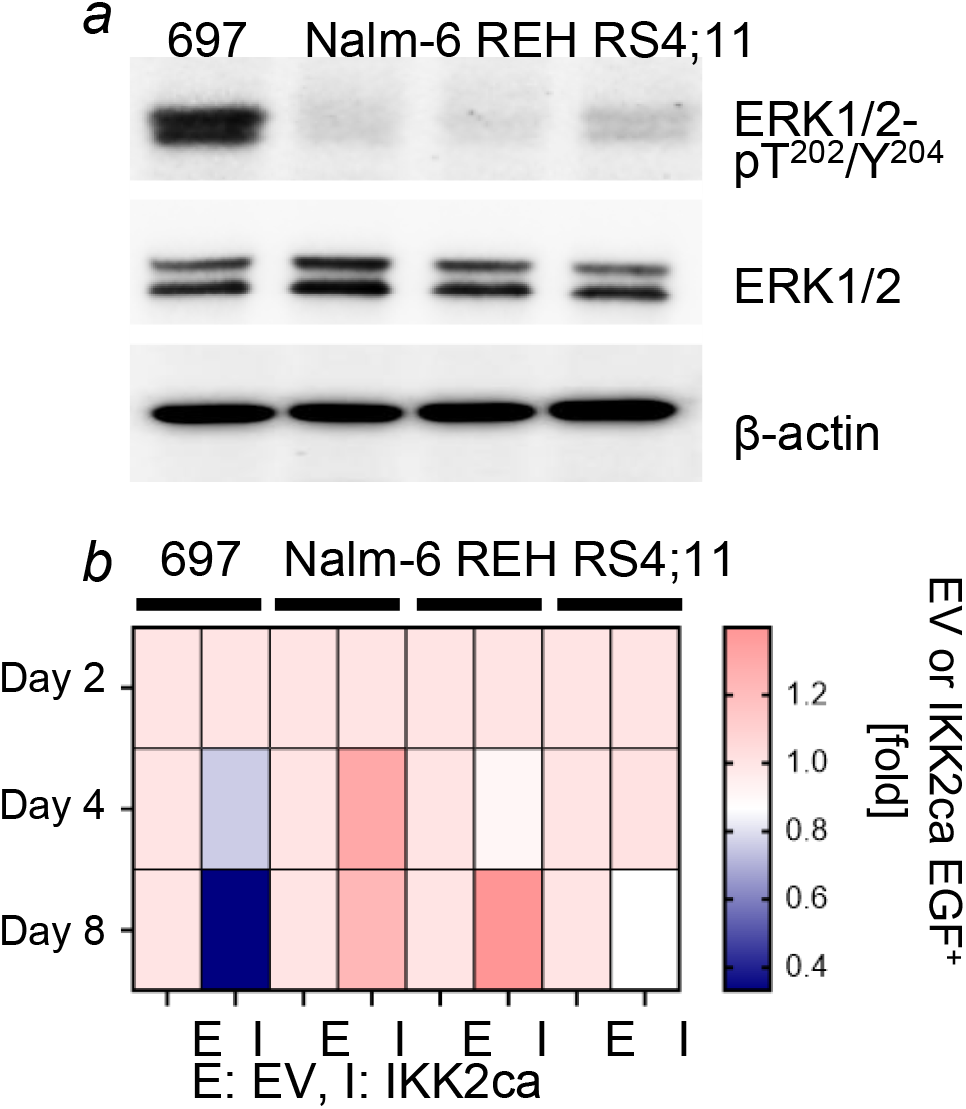
ERK activity determines the outcome of NF-κB activation in human B-ALL cells. (***a***) Western blots of human B-ALL cell lines. (***b***) Human B-ALL cell lines were transfected with empty vector (EV) or IKK2ca-EGFP. Cells were transfected using electroporation. Enrichment or depletion of EGFP+^+^ cells was monitored by flow cytometry at the indicated time points. Shown are mean values from 3 independent experiments.

To test whether pharmacologic activation of NF-κB produced similar effects, we stimulated Toll-like receptor (TLR) signaling, which activates NF-κB through the MYD88 adaptor pathway. Treatment with the TLR1/2 agonist CU-T12-9^45^ increased phosphorylation of NF-κB p65 while concurrently reducing ERK1/2 phosphorylation in RAS-pathway–mutated B-ALL cells (**Fig. 8a**). Treatment with either CU-T12-9 or an ERK inhibitor alone reduced cellular viability, and combined treatment produced a further decrease (**Fig. 8b**). These effects were selective for RAS-pathway–mutated contexts, as cell lines lacking RAS-pathway alterations showed minimal response (**Fig. 8c**).

**Figure 8:**
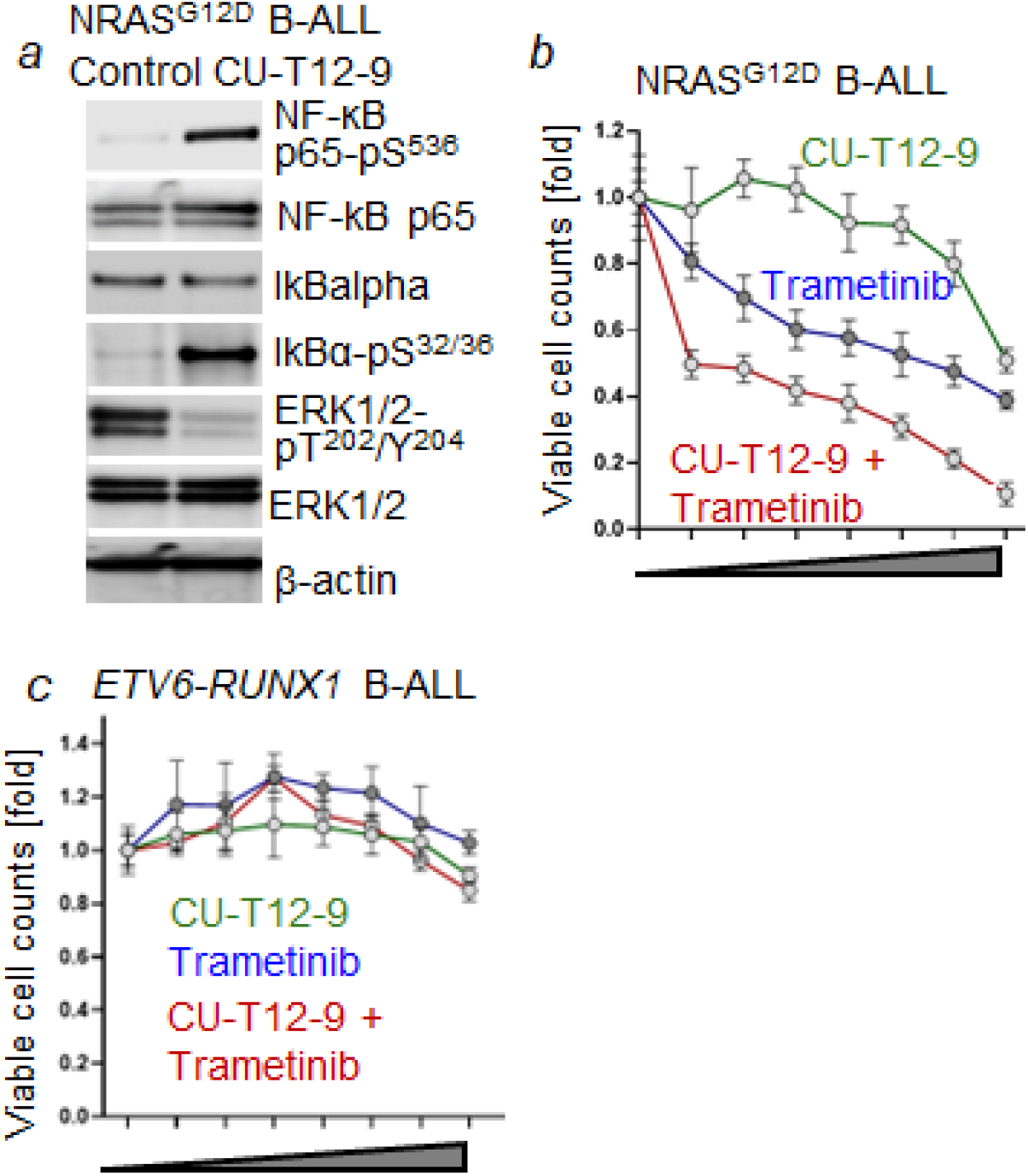
Incompatibility between NF-κB- and RAS-pathways as a therapeutic target in B-ALL. (***a***) Western blots of human NRAS^G12D^ B-ALL cells following treatment with control or CU-T12-9. (***b-c***) Human NRAS^G12D^ B-ALL (***b***) and *ETV6-RUNX1* B-ALL (no RAS-pathway mutations, ***c***) cells were treated with vehicle control, CU-T12-9 alone (green line), trametinib (blue line) alone, a combination of both (red line) for 72 hr. Viable cell counts (n=3) were determined. Shown are mean values of fold change relative to vehicle control (n=3). CUT129 (uM): 0, 0.156, 0.313, 0.625, 1.25, 2.5, 5, 10; Trametinib (nM): 0. 3.9, 7.8, 15.6, 31.3, 62.5, 125, 250.

Although frontline regimens combining glucocorticoids such as prednisolone with cytotoxic agents including vincristine achieve high remission rates, activating lesions in the RAS pathway are associated with therapeutic resistance and relapse, where survival rates decline to approximately 50%^8–10,46,47.^ The antagonistic interaction between NF-κB and RAS–ERK signaling suggests that activation of NF-κB can suppress ERK signaling and impair the viability of RAS-pathway–mutated B-ALL cells. Consistent with the concept of pathway incompatibility described previously for other signaling pairs^11^, pharmacologic activation of NF-κB in combination with ERK inhibition further reduced leukemic cell viability, supporting the broader principle that enforcement of incompatible signaling states can constrain oncogenic signaling programs.

## DISCUSSION

Oncogenic mutations are commonly interpreted as cooperating to promote malignant transformation, and mutually exclusive alterations observed across cancers are often attributed to functional redundancy. The findings presented here support an alternative model in B-ALL in which specific signaling programs are incompatible and constrain leukemic fitness. Our data demonstrate that canonical NF-κB signaling antagonizes RAS–ERK activity and suppresses RAS-driven leukemogenesis, indicating that cellular signaling context determines whether oncogenic pathways reinforce or oppose malignant growth.

Genomic interaction analysis revealed marked underrepresentation of tumors harboring concurrent RAS and NF-κB pathway alterations, suggesting negative selective pressure against their coexistence. Functional experiments supported this genetic relationship: activation of canonical NF-κB induced apoptotic depletion of RAS-mutant leukemic cells, whereas inhibition of NF-κB signaling conferred a competitive growth advantage. In contrast, activation of the non-canonical pathway similarly conferred a growth advantage in competition assays, highlighting functional divergence between canonical and non-canonical NF-κB signaling.

Mechanistically, NF-κB–mediated suppression of RAS-driven leukemic fitness was associated with coordinated disruption of pre-BCR–associated signaling programs. Canonical NF-κB activation reduced expression of Bcl6, a critical effector previously shown to mediate pre-BCR–like signaling required for survival and proliferation of RAS-driven B-ALL cells^11^. This effect was accompanied by increased inhibitory phosphorylation of Foxo1 and reduced phosphorylation of the receptor-proximal kinase Syk, indicating attenuation of both transcriptional and signaling components of the pre-BCR network. Concurrently, NF-κB activation increased expression of mature BCR components, consistent with a shift in receptor composition away from pre-BCR–associated signaling. Because oncogenic RAS functionally mimics pre-BCR signaling to sustain leukemic survival^11^, disruption of this receptor-dependent program provides a mechanistic explanation for the observed loss of cellular fitness.

Consistent with this model, activating RAS mutations were enriched in malignancies arising from developmental stages lacking surface BCR expression^40^, and enforced expression of oncogenic RAS was poorly tolerated in BCR-positive lymphoma cells unless receptor expression was disrupted. These findings indicate that receptor-defined developmental state constrains the compatibility of oncogenic signaling programs and help explain why RAS activation preferentially supports transformation within specific stages of B-cell differentiation.

The functional consequences of this incompatibility suggest potential translational relevance. Canonical NF-κB activation reduced ERK signaling and selectively impaired viability of RAS-pathway mutated B-ALL cells, with enhanced effects observed in cells exhibiting high baseline ERK activity. Pharmacologic activation of NF-κB similarly suppressed ERK signaling and cooperated with ERK inhibition to reduce leukemic cell viability, supporting the concept that enforcement of incompatible signaling states can selectively constrain RAS-dependent leukemogenesis.

Several questions remain. The molecular intermediates linking NF-κB activation to attenuation of ERK signaling and suppression of pre-BCR–associated programs remain incompletely defined and may involve transcriptional or chromatin-level regulation downstream of NF-κB. In addition, validation across genetically diverse leukemic contexts and *in vivo* models will be necessary to fully define the scope and therapeutic implications of this incompatibility. Whether similar constraints operate between other oncogenic signaling pathways also warrants investigation.

In summary, our findings demonstrate that canonical NF-κB signaling restricts RAS-driven leukemogenesis by suppressing pre-BCR–dependent survival programs and enforcing a receptor context incompatible with oncogenic RAS activity. These results provide mechanistic support for pathway incompatibility as a constraint on the oncogenicity of signaling pathways and identify developmental signaling state as a key determinant of leukemogenic potential.

## METHODS

### Human leukemia and lymphoma cell lines

Human leukemia and lymphoma cells used in this study were listed in **Table S1** and maintained in Roswell Park Memorial Institute medium (RPMI-1640; Invitrogen, Carlsbad, CA) with GlutaMAX containing 10%-20% FBS, 100 IU/mL penicillin and 100 μg/mL streptomycin. Cell cultures were kept at 37°C in a humidified incubator with 5% CO2.

### Extraction of bone marrow cells from mice

Bone marrow cells from 8-12 weeks old mice (**Table S2**) were harvested by flushing cavities of femur and tibia with chilled PBS followed by filtering through 40 μm strainer to yield a single-cell suspension. Filtered cells were further incubated with lysis buffer (RBC Lysis Buffer, BioLegend) to lyse erythrocytes. After washing with PBS, cells were subjected to further experiments. All mouse experiments were subject to approval by the Institutional Animal Care and Use Committee of the Cleveland Clinic Research.

### Murine model of B-cell acute lymphoblastic leukemia

Bone marrow cells were harvested and cultured in Iscove’s modified Dulbecco’s medium (IMDM; GIBCO) with GlutaMAX containing 20% FBS, 50 μM 2-mercaptoethanol, 100 IU ml^-1^ penicillin, 100 μg ml^-1^ streptomycin in the presence of 10 ng ml^-1^ recombinant mouse IL-7 (Peprotech). After one week of *in vitro* culture in the presence of IL-7, cells were confirmed to be nearly 100% Cd19 positive by flow cytometry. For the leukemia model, B cell precursor cells were retrovirally transduced with NRAS^G12D^-puromycin or NRAS^G12D^-K.orange followed by puromycin (1 µg/mL) selection or cell sorting for K.orange^+^ populations, respectively. Cells were propagated only for short periods of time and usually not longer than 2 months to avoid acquisition of additional genetic lesions during long-term cell culture.

## Retroviral transduction

Constructs used for retroviral transduction are listed in **Table S3**. Retroviral supernatant was produced by transient co-transfection of Lenti-X 293T cells with retroviral constructs together with packaging plasmid pHIT60 (gag-pol) and envelop plasmid pHIT123^11^ (gifts from Dr. Markus Müschen) using jetOPTIMUS. Triple-constructs transfected Lenti-X 293T cells were cultured on the poly-lysine-coated plate in high glucose Dulbecco’s Modified Eagle’s Medium (DMEM, GIBCO) with GlutaMAX containing 10% fetal bovine serum, 100 μg/mL streptomycin, 100 IU/mL penicillin, 1 mM sodium pyruvate and 0.1 mM non-essential amino acids for 16 hours. To boost virus production, sodium butyrate (10 mM) was added to the culture for an additional 8 hours and then washed out. The virus supernatant was collected 18-24 hours after sodium butyrate induction and filtered through a 0.45 μm filter. For retroviral transduction, the non-tissue culture-treated plate was coated with 50 μg/mL Retronectin (Takara), and the virus-containing supernatants were loaded by centrifugation (2,000 g, 90 min at 32ºC). Two to three million cells per well were loaded on the plate by centrifugation at 600 g for 30 min in the appropriate medium and cultured at 37ºC for 48 hr. Cells transduced with constructs carrying an antibiotic resistance marker were selected with its respective antibiotic.

### shRNA knockdown of Nfkb1

Lentiviral constructs carrying GFP-tagged shRNAs targeting *Nfkb1* or non-targeting control shRNA were purchased from Transomic Technologies. Lentivirus production and transductions were performed as described above, followed by cell sorting to select for cells expressing shRNA. Knockdown efficiency was verified by Western blot.

### Electroporation-based transfection

Transfection of 2 million human cells with plasmids (5-10 µg) encoding genes of interest or empty vector (EV) controls (**Table S3**) was performed using the Neon NxT Transfection System (Invitrogen). Cells were resuspended in a Neon NxT buffer (100 µL) and subjected to electrical pulses to introduce plasmid DNA (Pulse: 1600V, width: 10 ms, number of pulses: 3 pulses). Following transfection, cells were returned to standard culture conditions for recovery. Cells were analyzed by flow cytometry 48–72 hours post-transfection. Plasmids were purchased from VectorBuilder.

### CRISPR-mediated gene deletion

For non-viral gene deletion of conventional kappa light chains (Ig kappa LC, *IGKC*) in human cells, Alt-R CRISPR-Cas9 guide RNAs (**Table S4**) and non-targeting control guide RNAs (Cat # 1072544, 1072545, 1072546) were purchased from IDT. Chemically synthesized crRNAs (100 µmol/L) and tracrRNAs (100 µmol/L) were annealed by incubation at 95 Cº for 5 min. Recombinantly produced Cas9 (40 µmol/L) were then added to RNA mixture to produce RNA ribonucleoprotein (RNP) complexes. Electroporation was performed by using the Neon NxT Transfection system (Invitrogen). Non-targeting control crRNAs were purchased from IDT.

### Western blotting

Cells were lysed in CelLytic buffer (Sigma-Aldrich) supplemented with 1% protease inhibitor cocktail (Roche Diagnostics), 1% phosphatase inhibitor cocktail (EMD Millipore) and 1mM PMSF on ice. 15 - 20 µg of cell lysates per sample were separated on mini precast gels (Bio-Rad) and transferred on nitrocellulose membranes (Bio-Rad). Membranes were probed with the appropriate primary antibodies listed in **Table S5**. Membranes were incubated with alkaline phosphatase conjugated secondary antibodies (Invitrogen) and chemiluminescent substrate (Invitrogen) and were further detected by film exposure or ChemiDocTM MP Imaging System (Bio-Rad).

### Flow cytometry

Approximately 10^6^ cells per sample were resuspended in PBS blocked using Fc blocker for 10 minutes on ice, followed by staining with the appropriate dilution of the antibodies or their respective isotype controls for 15 minutes on ice. Cells were washed and resuspended in PBS with SYTOX® Blue stain (1 µM) as a dead cell marker. The antibodies used for flow cytometry are listed in **Table S5**. For competitive growth assays, percentage of GFP^+^, K.orange^+^, mCherry+ cells was monitored by a LSRFortessa™ flow cytometer (BD Biosciences). For Annexin V staining (detection of apoptosis), Annexin V binding buffer (BD Bioscience) was used instead of PBS and SYTOX® Blue stain (Thermo Fisher Scientific) was used as a dead cell marker. PE labeled-Annexin V was purchased from Thermo Fisher Scientific FACS data were analyzed with FlowJo software (FlowJo, LLC).

### Cell viability assays

Forty thousand patient-derived B-ALL cells were seeded in a volume of 80 μL in complete growth medium on 96-well plates (BD Biosciences). Compounds were added at the indicated concentration in a total volume of 100 μL. After culturing for 3 days, Cell Titer Glo (Promega) assays were performed according to the manufacturer’s instructions. Medium without cells was used as blank. Relative viability was calculated using baseline values of cells treated with vehicle control as a reference.

### Statistical Analyses

Data are presented as mean ± standard deviation (SD) from three independent experiments unless otherwise indicated. Statistical significance was determined using Student’s *t*-test for comparisons between two groups and one-way ANOVA for comparisons among three or more groups, with *P* < 0.05 considered significant. Analyses were performed using GraphPad Prism.

## ACKNOWLEDGEMENT

This research was supported in part by Cleveland Clinic Research Flow Cytometry Shared Laboratory Resource (Cleveland, OH) (RRID:SCR_026460). This work was supported by a DeLuca Center for Innovation in Hematology Research Pilot Grant (LNC), a grant from the When Everyone Survives (WES) Leukemia Foundation (LNC), a VeloSano 10 Pediatric Pilot Award (LNC) and an Ohio Cancer Research Grant (LNC).

## SUPPLEMENTARY TABLES

**Table S1.** Overview of lymphoma cell line used in this study.

| Cell line | Cancer type | Source | ID | Additional information |
| --- | --- | --- | --- | --- |
| 697 | B-ALL, childhood | Creative Bioarray | CSC-C0217 | NRAS<sup>G12D</sup>; expression of fusion gene *TCF3-PBX (E2A-PBX)* |
| RS4;11 | B-ALL, adult | ATCC | CRL-1873 | *MLL-AF4 (KMT2A-AFF1)* |
| Nalm-6 | B-ALL, adult | ATCC | CRL-3273 | NRAS<sup>A146T</sup>; *ETV6-PDGFRB* |
| REH | B-ALL, childhood | ATCC | CRL-8286 | *ETV6-RUNX1* |
| U2932 | DLBCL | Creative Bioarray | CSC-C0624 |  |
| TMD8 | DLBCL | Dr. Neetu Gupta, CCF |  | MYD88; p.Leu252Pro (c.755T>C) (L265P) |
| OCILY3 | DLBCL | Dr. Neetu Gupta, CCF |  | MYD88; p.Leu252Pro (c.755T>C) (L265P) |

**Notes:** B-ALL: B-cell acute lymphoblastic leukemia; DLBCL: Diffuse large B-cell lymphoma; ATCC: American Type Culture Collection; CCF: Cleveland Clinic Foundation

**Table S2.**
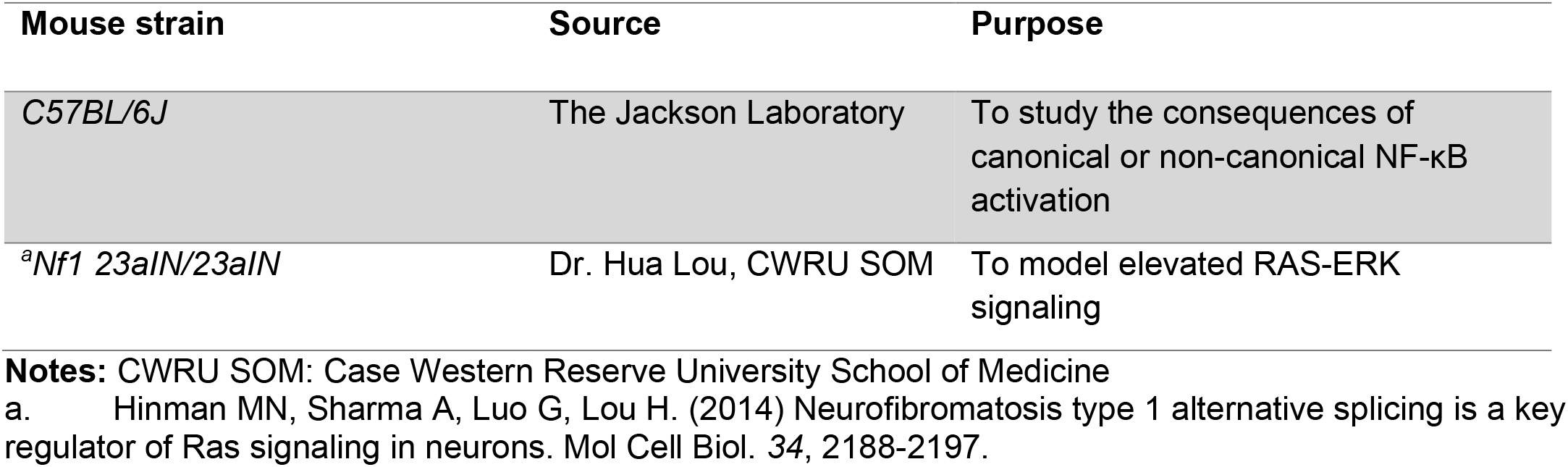
Overview of genetic mouse models used in this study.

| Mouse strain | Source | Purpose |
| --- | --- | --- |
| C57BL/6J | The Jackson Laboratory | To study the consequences of canonical or non-canonical NF-κB activation |
| <sup>a</sup>Nf1 23aIN/23aIN | Dr. Hua Lou, CWRU SOM | To model elevated RAS-ERK signaling |

**Notes:** CWRU SOM: Case Western Reserve University School of Medicine
a. Hinman MN, Sharma A, Luo G, Lou H. (2014) Neurofibromatosis type 1 alternative splicing is a key regulator of Ras signaling in neurons. Mol Cell Biol. 34, 2188-2197.

**Table S3.** Mammalian gene expression vectors -regular plasmids and retrovirus constructs.

| Construct | Expression of | Purpose |
| --- | --- | --- |
| MSCV NRAS<sup>G12D</sup>-Puro | NRAS<sup>G12D</sup>; Puromycin resistance | Transformation; RAS-activation (retroviral transduction) |
| pCGDNSam-NRAS<sup>G12D</sup>-IRES-Kusabira orange | NRAS<sup>G12D</sup>; Kusabira orange | Transformation; RAS-activation (retroviral transduction) |
| pMIG EV | GFP | Empty vector control (retroviral transduction) |
| pMSCV IKBKB S177E/S181E EGFP | Ikk2ca mutant, EGFP | Canonical NF-κB activation (retroviral transduction) |
| pMSCV MAP3K14 (NIK) EGFP | Wild-type NIK, EGFP | Noncanonical NF-κB activation (retroviral transduction) |
| pMIG IKBSR | Super-repressor form of IκBα, GFP | Modulation of NF-κB activity |
| pRP CMV EGFP | EGFP | Empty vector control (transfection) |
| pRP EGFP CMV IKBKB S177E/S181E | Ikk2ca mutant, EGFP | Canonical NF-κB activation (transfection) |
| pRP CMV mCherry | mCherry | Empty vector control (transfection) |
| pRP mCherry CMV NRAS<sup>G12D</sup> | NRAS<sup>G12D</sup>; mCherry | Transformation; RAS-activation (transfection) |

**Notes:** NRAS<sup>G12D</sup> constructs: gifts from Dr. Markus Müschen; other constructs were purchased from VectorBuilder

**Table S4.**
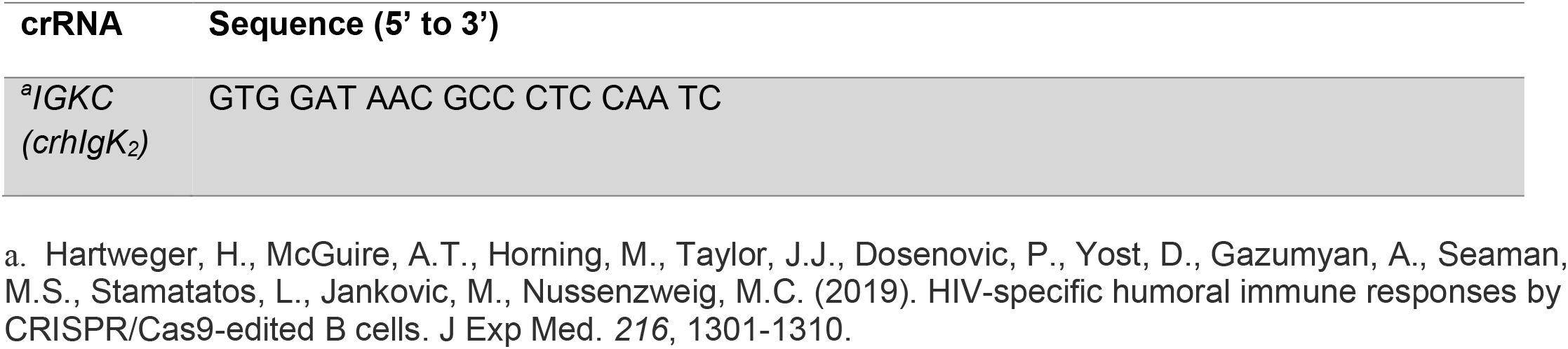
Oligonucleotide sequences.

**Table S5.** Antibodies used in this study.

| **Antigen** | **ID** | **Manufacture** |
| --- | --- | --- |
| ERK-pT<sup>202</sup>/Y<sup>204</sup> | 4370 | Cell Signaling Technology |
| ERK | 9102 | Cell Signaling Technology |
| β-actin | 3700 | Cell Signaling Technology |
| NF-κB1 p105/p50 | 13586 | Cell Signaling Technology |
| NF-κB p65-pS<sup>536</sup> | 3033 | Cell Signaling Technology |
| NF-κB p65 | 8242 | Cell Signaling Technology |
| BCL6 | 5650 | Cell Signaling Technology |
| Zap-70 pY<sup>319</sup> Syk p<sup>Y352</sup> | 2717 | Cell Signaling Technology |
| Zap-70 | 3165 | Cell Signaling Technology |
| Syk | 13198 | Cell Signaling Technology |
| FoxO1 pS<sup>256</sup> | 9461 | Cell Signaling Technology |
| FoxO1 | 2880 | Cell Signaling Technology |
| IκBα pS<sup>32/36</sup> | 9246 | Cell Signaling Technology |
| IκBα | 9242 | Cell Signaling Technology |

*Flow cytometry antibodies for mouse cells*
| **Antigen** | **Clone ID** | **Manufacturer** |
| --- | --- | --- |
| TruStain fcX Fc block |  | Biolegend |
| Ig light chain kappa | 187.1 | D Biosciences |
| Ig light chain lambda | RML-42; R26-46 | Biolegend, BD Biosciences |
| IgM | RMM-1 | Biolegend |

*Flow cytometry antibody for human cells*
| **Antigen** | **ID** | **Manufacturer** |
| --- | --- | --- |
| Ig κ light chain | TB28-2 | Biolegend; BD Biosciences |
| Ig λ light chain | MHL-38 | Biolegend |
| TruStain fcX Fc block |  | Biolegend |

